# Placing approach-avoidance conflict within the framework of multi-objective reinforcement learning

**DOI:** 10.1101/2023.01.05.522878

**Authors:** Enkhzaya Enkhtaivan, Joel Nishimura, Amy Cochran

## Abstract

Many psychiatric disorders are marked by impaired decision-making during an approach-avoidance conflict. Current experiments elicit approach-avoidance conflicts in bandit tasks by pairing an individual’s actions with consequences that are simultaneously desirable (reward) and undesirable (harm). We frame approach-avoidance conflict tasks as a multi-objective multi-armed bandit. By defining a general decision-maker as a limiting sequence of actions, we disentangle the decision process from learning. Each decision maker can then be identified as a multi-dimensional point representing its long-term average expected outcomes, while different decision making models can be associated by the geometry of their ‘feasible region’, the set of all possible long term performances on a fixed task. We introduce three example decision-makers based on popular reinforcement learning models and characterize their feasible regions, including whether they can be Pareto optimal. From this perspective, we find that existing tasks are unable to distinguish between the three examples of decision-makers. We show how to design new tasks whose geometric structure can be used to better distinguish between decision-makers. These findings are expected to guide the design of approach-avoidance conflict tasks and the modeling of resulting decision-making behavior.

## 1 Introduction

Approach-avoidance conflict (AAC) refers to a scenario where a person has an experience or a goal that can simultaneously have both desirable and undesirable consequences (Bach et al., 2014). For example, someone may be deciding between taking a new job that pays better than their current job (*approach*) or staying at their current job because it requires working less hours (*avoid*). Researchers want to understand how different individuals make decisions and learn during an AAC (Epstein & Fenz, 1962; Miller, Marcotulli, Shen, & Zweifel, 2019; Zorowitz et al., 2019), since AAC is implicated in a number of psychiatric disorders, such as social anxiety disorder (Pittig, Brand, Pawlikowski, & Alpers, 2014) or post-traumatic stress disorder (Weaver et al.,2020). The overarching goal is to identify mechanisms by which one person might forgo desirable consequences in an effort to avoid undesirable ones.

A common strategy is to devise an experiment in which individuals have to make decisions sequentially over time during an AAC (Aupperle, Sullivan, Melrose, Paulus, & Stein, 2011; Bach et al., 2014; Pittig et al., 2014; Weaver et al., 2020; Zorowitz, Momennejad, & Daw, 2020). The decision-making scenario in these experiments can be described as follows: on trial *t*, a person is in a current state *S_t_* and takes an action *A_t_*, after which there is a consequence *R_t_* and a transition to a possibly new state *S*_*t*+1_. Such an experiment would be similar to other sequential-decision making experiments used in psychology except for the following difference: *R_t_* could be multi-dimensional in order to capture the simultaneous occurrence of multiple consequences. Some consequences are desirable (e.g., monetary reward) and some are not (e.g., an electric shock or a threatening image). The researcher analyzes the actions *A*_1_,…,*A_t_* that an individual makes in the face of conflicting consequences.

Actions in sequential decision-making experiments are analyzed with the help of temporal-difference (TD) learning models, which were adopted from the reinforcement learning literature to explain human decision-making. A TD learning model consists of two processes: a learning process which updates how the agent values a decision and a decision-making process which determines how the agent selects an action based on their current valuation (Sutton & Barto, 2018). One popular TD learning model is called Q-learning. It is described by the update equations:

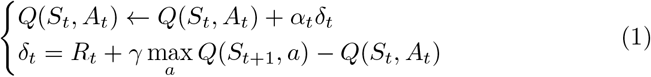

where *Q*(*s, a*) is the learned value placed on action *a* in state *s*, *α_t_* is the learning rate parameter, and *δ_t_* is a prediction error in expected future rewards discounted by *γ*. The person would then weigh different actions in state *S_t_* according to *Q*(*S_t_*, ·). An important connection between TD learning and human learning was made by Schultz, Dayan, and Montague (1997), who proposed that dopamine fluctuations in the brain encodes information about prediction errors between expected reward and actual reward. This hypothesis provides a biological explanation for how prediction error, *δ_t_*, guides how a human learns about the environment. Since this seminal work, reinforcement learning models have been adapted in order to describe different aspects of human decision-making such as sensitivity to risk (Enkhtaivan, Nishimura, Ly, & Cochran, 2021; Niv, Edlund, Dayan, & O’ Doherty, 2012; Ross, Lenow, Kilts, & Cisler, 2018) or learning latent states (Cochran & Cisler, 2019; Gershman, Blei, & Niv, 2010; Redish, Jensen, Johnson, & Kurth-Nelson, 2007). The field would similarly benefit from a TD learning model to describe human decision-making during an AAC with multi-dimensional outcomes *R_t_*.

Adapting a TD learning model to describe decision-making during an AAC requires strategies for handling the multi-dimensionality of *R_t_*. Multidimensional *R_t_* can be handled for the learning process by simply defining update equations like (1) to update a *Q* value along each dimension of *R_t_*, resulting in a table of *Q* values. While such will learn, by itself it is unable to negotiate tradeoffs between different objectives. Instead multiple objectives requires a decision making model to weigh different actions according to the table of *Q* values. As a starting point, we look to the emerging frameworks of multi-objective reinforcement learning, dedicated to solving reinforcement learning problems that involve optimizing a multidimensional objective. In particular, we will use the multi-objective multi-armed bandit (MO-MAB) framework of Drugan and Nowe (2013) since it’s common to model human decision-making experiments using a multi-armed bandit model (Reverdy, Srivastava, & Leonard, 2014; Steyvers, Lee, & Wagenmakers, 2009).

What makes a MO-MAB problem more difficult than its single-objective counterpart is that two actions can outperform each other along different dimensions of the objective function. Such a conflict is frequently handled in one of two ways. The first is *scalarization*, which transforms a MO-MAB problem into a standard, single-objective bandit problem. The second is to weaken the concept of optimality to *Pareto optimality*, whereby an action is Pareto optimal if it cannot be out-performed by another action along every dimension, and search for Pareto optimal solutions. To date, MO-MAB, or more generally, multi-objective reinforcement learning, is largely focused on: obtaining (Pareto) optimal solutions, or near-optimal solutions; algorithms for solving industrial tasks such as water-reservoir control (Castelletti, Corani, Rizzolli, Soncinie-Sessa, & Weber, 2002); the management of energy consumption (Kwak et al., 2012); and optimizing broker agents that maximize profit and minimize spreads (Shelton, 2001). In contrast, relatively little work has been used the MO-MAB framework to understand human decision-making.

The goal of this paper is to view human decision-making during AAC through the lens of a MO-MAB framework, in an effort to guide the design and modeling of sequential decision making tasks for studying AAC. We want to answer the following questions:

1. If we fix a sequential decision-making task, what type of behavior do we observe during AAC when different models of decision-makers are utilized?
2. If we allow the task to vary, what tasks are better suited for distinguishing between different models of decision-makers?

To simplify what follows, we focus on the decision-making process of human decision-making, as opposed to the learning process, and on bandit tasks for which the state *S_t_* is constant (i.e. serial actions have independent consequences).

The organization of the paper is as follows. In Section 2, we introduce MO-MAB more formally. In Section 3, we use scalarization to introduce multi-objective versions of popular decision-makers in computational psychiatry that rely on softmax and epsilon-greedy rules. We also introduce a third decision-maker as an example of non-scalarized approach and show that this decision-maker can yield different qualitative behavior than the other two decision-makers. In Section 4, we show that every possible decision-maker in prior AAC tasks (Aupperle et al., 2011; Pittig et al., 2014; Weaver et al., 2020) is Pareto optimal. In Section 5, we introduce tasks for which these TD learning models do exhibit very different qualitative behaviors. Furthermore, we show that qualitatively different behavior between TD learning models can be revealed as a result of a small perturbation to the structure of the task. Last, we discuss our findings in the context of the literature in Section 6.

## 2 Multi-objective multi-armed bandits

We begin our investigation by formally introducing the MO-MAB framework from Drugan and Nowe (2013). MO-MAB draws from two areas. The first area is a special case of reinforcement learning problems known as multi-armed bandits. The second area is multi-objective optimization, which extends familiar optimization problems to ones in which multiple, rather than single, objectives are optimized.

### 2.1 Multi-armed bandits

Many sequential decision-making tasks in psychology are described by a multi-armed bandit problem of the following form. An individual has to make sequential decisions under uncertainty. Each action brings rewards ex ante unknown to the individual. The goal of the individual is to maximize the total accumulated reward. Formally, let 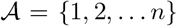 be the set of all possible actions and the individual selects an action 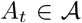 on trial *t*, (*t* = 1, 2,…). If action *i* is selected on trial *t*, then the individual receives an independent, random reward *R_t_* with finite mean *μ_i_*. An instance of a multi-armed bandit problem would then look like:

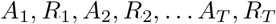

if a total of *T* rounds are played.

Traditionally, each reward would be scalar-valued, in which case this decision-making scenario is called a *single-objective* multi-armed bandit. We emphasize that rewards depend only the action taken and not some state of the system, which makes a multi-armed bandit problem a special case of a reinforcement learning problem. Furthermore, we note here that *A_t_* is a random variable taking values in 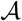 and can only use information from past actions and rewards up to time *t*. Formally, we take this latter assumption to mean that the *σ*-algebra generated by *A*_1_, *R*_1_,…,*A_t_*, *R_t_* is a subset of the *σ*-algebra generated by *A*_1_, *R*_1_,…,*A*_*t*+1_, *R*_*t*+1_ for all *t* ≥ 1.

Studying AAC requires at least two objectives, one which has desirable consequences (reward) and one which has undesirable consequences (harm). For simplicity, we assume rewards and harm can be encoded numerically as a continuous or binary variable such that increasing values are more desirable. A sequential decision-making experiment for AAC would thus be described by *multi-objective* multi-armed bandit problems. These are identical to their single-objective counterparts except now the reward is vector-valued, rather than scalar-valued. If action *i* is selected on trial *t*, then ***R**_t_* is an independent random vector sampled from a distribution *F* with vector-valued mean ***μ**_i_*.

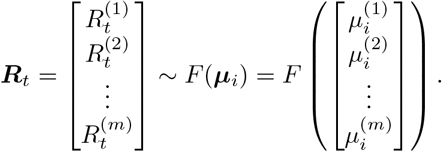

The goal of the individual is to maximize the total accumulated reward *along every dimension*.

An AAC arises when each action yields different vector-valued means: ***μ***_1_,…,***μ**_n_*. We thus characterize the underlying reward structure of an AAC task with *m* outcome dimensions and *n* possible actions as the matrix:

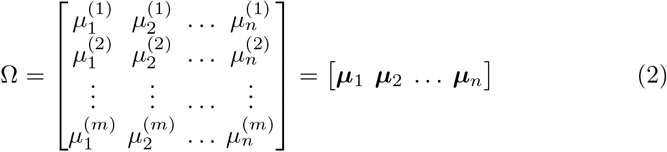

For example, the three AAC tasks we will study in Section 4 will be described by a 2-by-*n* matrix Ω capturing AAC with two outcome dimensions. Lastly, for our analysis of models and tasks for AAC in sections 2 and 5, we can avoid certain pathological cases by making the simplifying assumption that Ω is not *degenerate* in the following sense:

**Assumption 1** The convex hull of Ω, viewed as a convex polytope in 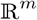, has the property that any *j*-dimensional face on its boundary contains exactly *j* points from {***μ***_1_, ***μ***_2_,…,**μ*_n_*}. Further, for any outcome dimension *j*, the components 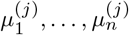 are pairwise distinct.

This assumption, for instance, ensures that the ***μ**_j_* on the boundary of the convex hull of Ω are pairwise distinct. It also ensures there is exactly one action that maximizes expected rewards along dimension *j*. Despite this simplifying assumption, the remaining possible Ω will be sufficiently expressive to capture all AAC tasks of practical interest to us. Indeed, while Assumption 1 is not true for all Ω, it is dense in the space of Ω, such that there is always some Ω′nearby for which Assumption 1 is true.

### 2.2 Multi-objective optimization

Since there may not be an action that maximizes expected reward along every dimension in a MO-MAB problem, we need to turn to multi-objective optimization to evaluate how well an individual makes decisions under conflicting objectives. A multi-objective optimization problem aims to optimize a vector function for which each entry represents an objective. For example, a maximization problem looks like:

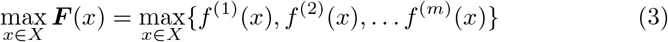

with *f*^(*i*)^(*x*) being the value of the *i*^th^ objective. In a non-trivial setting, this problem does not have a single solution *x* that simultaneously maximizes each *f*^(*i*)^(*x*); therefore, certain trade-offs between conflicting objectives have to be considered.

A popular way to describe optimality for multi-objective optimization is to use Pareto dominance. A solution *x* is said to *Pareto dominate* a solution *y* if:

1. *f*^(*i*)^(*x*) ≥ *f*^(*i*)^(*y*) for all *i* ∈ {1, 2,…*m*}
2. *f*^(*i*)^(*x*) > *f*^(*i*)^(*y*) for at least one *j* ∈ {1, 2,…*m*}

A solution *x* ∈ *X* is said to be *Pareto optimal* if it is Pareto dominated by no other solution *y* ∈ *X*. The set of all such x is called the *Pareto front* and finding it is often the ultimate goal of the optimization problem at (3).

A MO-MAB probem becomes a multi-objective optimization problem when wanting to maximize the long-term average of expected reward signals. Specifically, suppose the vector-valued rewards after *T* rounds received by the individual are

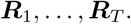

Total expected rewards accumulated by the individual would then be

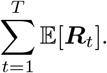

Note that the random variable ***R**_t_* draws from the reward distribution associated with arm i, when conditioned on the event *A_t_* = *i*. We are interested in the limiting behavior (should it exist):

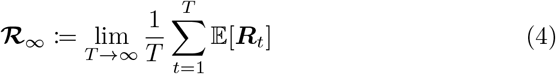

We can then say that an individual’s decision-making Pareto dominates another decision-making strategy for which the long-term average expected reward exists and is ***U***_∞_ if

1. 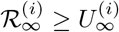 for all *i* ∈ {1, 2,…*m*}
2. 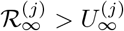 for at least one *j* ∈ {1, 2,…*m*}

## 3 Models of human decision-making

Within the context of MO-MAB, we can now introduce models that a researcher could use to investigate human decision-making during AAC. As noted earlier, we focus on the decision-making process as opposed to the learning process, or more specifically, on how a decision-maker performs with respect to long-term average expected rewards (Equation 4).

### 3.1 A general decision-maker

We want to consider decision-makers for whom this long-term average exists. A sufficient condition is that the actions *A_t_* converge in distribution to some random variable *A*_∞_. This motivates the following definition.

**Definition 1** Consider a MO-MAB problem with *m* objectives, *n* possible actions, and reward structure Ω defined in (2). We define a *decision-maker D* to be a sequence of actions on the action set 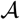:

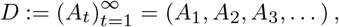

such that *A_t_* converges in distribution to a limiting random variable 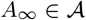. That is, if 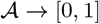 are the probability mass functions of *A_t_* and *A*_∞_, respectively, then:

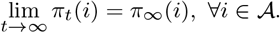

For a decision-maker *D*, one can quickly establish that the long-term average expected rewards obtained by *D* will be:

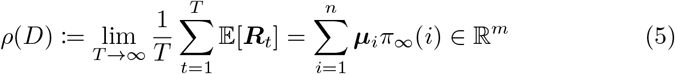

The fact that this converges is a simple consequence of Cesaro-Stolz theorem (see proof in Appendix A). It’s worth noting that our general decision-maker includes a decision-maker *D* who always selects action *i*:

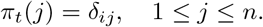

and whose long-term limit *ρ*(*D*) would be the vector ***μ**_i_*.

Let 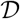 be the set of all decision-makers *D* defined as above. We will not only want to think about long-term average expected rewards *ρ*(*D*) for a single decision-maker *D* but also the possible long-term average expected rewards of a family 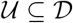 of decision-makers. This motivates the following:

**Definition 2** Given a collection of decision-makers 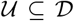, we define the *feasible region* of 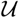 as:

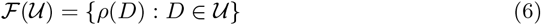

Notice that (5) tells us that the long-term average expected reward of a given decision-maker *D* is a weighted average of the reward vectors ***μ**_i_* that defines our AAC task. Intuitively then, if we consider the feasible region of all possible decision-makers, it should coincide with the convex hull of the ***μ**_i_*. In fact, we have the following proposition (its proof is given in Appendix B):

**Proposition 1** 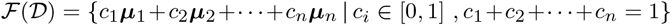

Let us consider an AAC task with a task structure described by

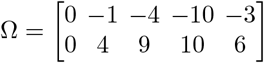

with action set 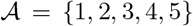. Here, action 1 yields rewards with mean [0 0]^*T*^, while action 4 yields rewards with mean [−10 10]^*T*^. The feasible region for this task is shaded in gray in Figure 1 and, as noted earlier, is exactly the convex hull of the ***μ**_i_*. Any decision-maker would yield long-term average expected costs somewhere in the gray region.

**Fig. 1.**
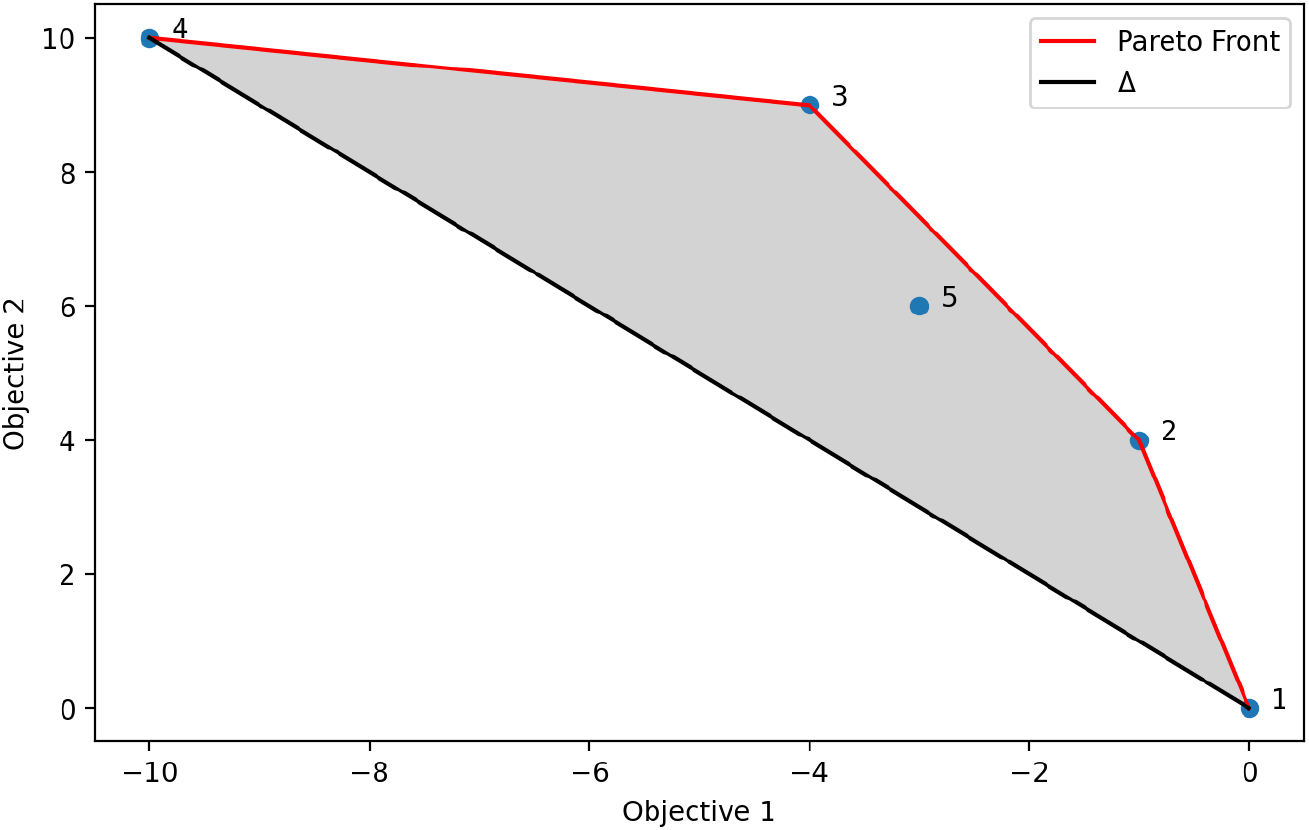
Feasible region of all decision-makers is shown in gray for an AAC task with two objectives to be maximized. The blue dots numbered 1 to 5 are the long-term average expected costs for the decision-makers that always selects the corresponding action. The Pareto front is shown in red. The line Δ connecting blue dot 1 and 4, each maximizing exactly one objective, is also shown.

Figure 1 shows two additional geometrical objects that will be of interest to us. The first is the Pareto Front 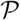 consisting of the long-term average expected rewards of any decision-maker that is not Pareto dominated by another decision-maker. The Pareto front in Figure 1 is indicated by the piecewise linear line segment in red. The second is Δ, the part of a hyperplane in 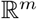 given by the convex hull of the (up to) *m* Pareto optimal points that maximize at least one of the objectives. For example, in the sketch above, Δ is the black line segment connecting points ***μ***_1_ and ***μ***_4_.

We point out that the Pareto front is not a subset of the vertices {***μ***_1_,…,***μ**_n_*}, as one might expect from the multi-objective optimization literature. This difference arises because decision-makers can mix actions. For instance, a decision-maker who chooses actions 4 and 3 each 50% of the time will be identified by the midpoint of vertices ***μ***_4_ and ***μ***_3_. Because of this, there might be a vertex ***μ**_i_* which is not Pareto dominated by any other vertices but is not part of the Pareto front 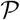. This is illustrated by vertex ***μ***_5_ in Figure 1, since the decision-maker who always selects action 5 is dominated by a decision-maker who mixes between action 2 and 3.

### 3.2 Three specific examples of decision-makers

We can introduce three specific examples of decision-makers by extending TD learning models to a MO-MAB setting. They represent different theories for the manner in which humans make decisions during an AAC.

#### 3.2.1 Learning

For a single-objective multi-armed bandit, a natural learning strategy for the individual would be to maintain an estimate *Q_t_*(*i*) for expected rewards of action *i* on trial *t* based on the rewards received so far. For example, if arm *i* was pulled for the *k*th time on trial *t* and the rewards received from this arm are *R*_*n*1_, *R*_*n*2_,…*R*_*n*__*k*_, then the individual could maintain the *sample-average*:

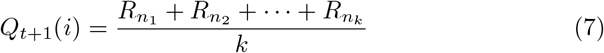

The strong law of large numbers tells us that if this arm is selected infinitely often in the limit *t* → ∞, then the estimate *Q*_t_(*i*) converges almost surely to the mean reward associated with said arm:

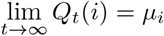

One can also write (7) equivalently as

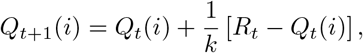

which is a special case of the TD learning model introduced at (1).

For a MO-MAB, the individual could learn about multi-dimensional rewards by maintaining an estimate 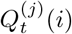 of expected rewards for each action *i and* each outcome dimension *j*. These estimates are then stored in a vector *Q_t_*(*i*). For example, if arm *i* was pulled for the *k*th time after trial *t* yielding rewards ***R**_t_* then the individual could update a table of *Q* values reflecting sample averages:

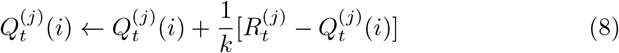

for each objective *j* = 1, 2,…*m*.

#### 3.2.2 Decision-making

With (7) and (8) dictating how an individual learns about rewards, let us now turn to how the individual makes a decision *A*_*t*+1_ based on:

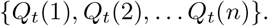

This is typically achieved by using this set to define a probability distribution *P*_*t*+1_ for selecting action *A*_*t*+1_ from 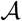. For example, if rewards were onedimensional, one can consider an epsilon-greedy decision-maker who, for a chosen parameter *ε* ∈ [0,1], samples uniformly from 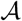 with probability *ε* and otherwise chooses the action with highest action-value estimate, *Q_t_*(·), with probability 1 − *ε*. That is, if we let

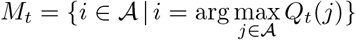

then:

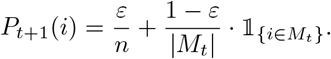

An important feature of the epsilon-greedy decision-maker is that it will eventually sample every arm infinitely often and thus can arrive at accurate estimates for each arm. An important drawback of the epsilon-greedy decision-maker is that the worst action is just as likely to be selected as the second-best action. This can be fixed by making *P*_*t*+1_(*i*) increase with increasing *Q_t_* values. With this in mind, one defines the softmax decision-maker for a parameter *τ* > 0 as someone who chooses action *A*_*t*+1_ with probability:

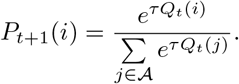

A softmax rule is a common choice in psychology for modeling human decision-making (Sutton & Barto, 2018).

When rewards are multi-dimensional, the issue now is how to decide which action to take since there are *m* different values of *Q* for each action. The most obvious choice is to consider a function:

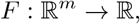

that transforms the m different values of *Q* into a scalar; this process is called *scalarization*. We can then make decisions based on the scalarized *Q* table:

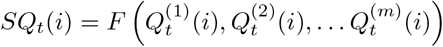

Consequently, this single value *SQ_t_*(*i*) can be used with typical action-selection strategies such as the epsilon-greedy or a softmax rule. As an example from the MO-MAB literature, the linear scalarization

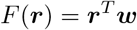

with a weight vector ***w*** = (*w*_1_, *w*_2_,…*w_m_*) that satisfies ∥**w**∥_1_ = *w*_1_ + *w*_2_+…+*w_m_* = 1, and *w_i_* ∈ [0, 1] was analyzed in Moffaert, Drugan, and Nowe (2013).

In light of these observations, we will consider decision-makers that apply an epsilon-greedy or a softmax rule to linear scalarization of the *Q* table, which follows the TD-learning rule as described in (8):

**Definition 3** Let ε ∈ [0,1], 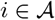, **w** ∈ [0, 1]^*m*^ with ∥**w**∥_1_ = 1 and define:

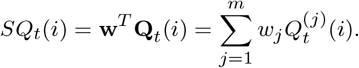

An *epsilon-greedy decision-maker* selects action *i* on trial *t* + 1 with probability:

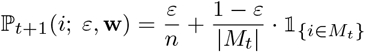

where

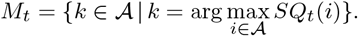

**Definition 4** Let *τ* ∈ [0, ∞), 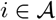, **w** ∈ [0,1]^*m*^ with ∥**w**∥_1_ = 1 and define:

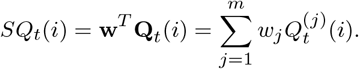

A *softmax decision-maker* selects action *i* on trial *t* + 1 with probability:

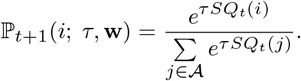

Since individuals might not resort to scalarization to make decisions, we provide an example of a decision-maker that is not scalarized. This decision-maker first restricts its attention to a specific reward dimension *j* with some probability *w_j_* and then makes the decision based on the *Q* values for the selected dimension using the softmax method. We note that this is based on the notion of selective attention, which purports that when there are multiple stimuli, people typically restrict their attentions to a few select stimuli (Johnston & Dark, 1986; Nishimura & Cochran, 2020; Treisman, 1969).

**Definition 5** Let *τ* ∈ [0, ∞), 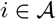, **w** ∈ [0,1]^*m*^ with ∥**w**∥_1_ = 1. A *selective attention decision-maker* selects action *i* on trial *t* + 1 with probability:

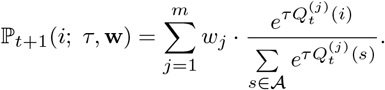

#### 3.2.3 Feasible region

While Definitions (3)–(5) are for fixed parameter values, we can consider a family of each type of decision-makers by letting these parameters range over all of their possible values. With this in mind, we were able to formally characterize properties of the feasible regions for the three families of decision-makers (leaving their proofs to the Appendix).

For epsilon greedy decision-makers we can exactly characterize the entire feasible region as:

**Proposition 2** For the family of epsilon-greedy decision-makers, the feasible region *is exactly* the set of line segments connecting the centroid of the vertices of Ω to each vertex on the Pareto front 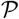 and the centroid of the vertices on each face on Pareto front 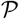.

For softmax decision-makers the feasible region is well understood by understanding it’s boundary, which is exactly given by:

**Proposition 3** For the family of softmax decision-maker, the feasible region *is bounded exactly by* the Pareto Front 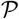 and the feasible region of a subset of softmax decision-makers for whom *w_j_* = 0 for one of the entries in parameter ***w***.

In comparison to epsilon-greedy and softmax decision makers, the feasible region of selective attention decision makers is harder to exactly characterize, especially in higher dimensions, but it’s boundary always includes two critical landmarks, the hyperplane Δ and the surfaces given by ***w**_j_* = 0:

**Proposition 4** For the family of selective attention decision-maker, the feasible region *is bounded by* the hyperplane Δ and the feasible region of the subset of selective attention decision-makers for whom *w_j_* = 0 for one of the entries in parameter ***w***

## 4 Previous AAC tasks

With our definition of decision-makers and their feasible regions, we are ready to analyze sequential decision-making experiments that have been previously used in the literature to study AAC. For each experiment, participants receive a reward (desirable consequence) and potential harm (undesirable stimulus) for each action they choose. While the rewards are numerical (either points or fictional money), potential harm can be electric shocks (Zorowitz et al., 2019) or disturbing visual and sound cues (Pittig et al., 2014; Weaver et al., 2020). We represent harm as a numerical value ranging from —1 for most harmful outcome to 0 for least harmful outcome (see Appendix D for a discussion on the degree to which this choice in numerical value matters); the corresponding objective is to maximize the numerical value. Table 1 summarizes reward structures Ω for each task during a phase or version of the experiment when there is an AAC.

### 4.1 Spider gambling task

Pittig et al. (2014) introduced the spider gambling task, which was based on the original Iowa gambling task of Bechara, Damasio, Damasio, and Anderson (1994). The Iowa gambilng task asks subjects to repeatedly select from one of four decks at a time, each of which has different reward distribution. The first two decks *A* and *B* (referred as bad decks), yielded net loss in the long term, while decks *C* and *D* (referred as good decks) yielded long term gains. The spider gambling task introduced another outcome dimension to the Iowa gambling task by adding pictures of spiders and butterflies to the good and bad decks, respectively, thus creating a conflict. There was also a non-conflict version where the pictures of spiders were on the back of the bad decks as well. The study was undertaken by participants that were afraid of spiders and control participants.

**Table 1.** Reward structure Ω from AAC tasks used in the literature

| AAC task | $\Omega$ |
| --- | --- |
| Spider gambling | $\begin{bmatrix} -1 & -1 & 0 & 0 \\ 0 & 0 & 50 & 50 \end{bmatrix}$ |
| Weather | $\begin{bmatrix} -1.0 & -0.875 & -0.75 & -0.625 & -0.5 & -0.375 & -0.25 & -0.125 & 0.0 \\ 1.8 & 1.6 & 1.4 & 1.2 & 1.0 & 0.8 & 0.6 & 0.4 & 0.2 \end{bmatrix}$ |
| Trauma-related AAC | $\begin{bmatrix} 0.0 & -0.5 & -1.0 \\ 0.2 & 0.5 & 0.8 \end{bmatrix}$ |

Figure 2 illustrates the feasible region 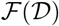 of all decision-makers and the the Pareto front for the spider gambling task. For both the conflict and non-conflict version, selecting deck *A* is indistinguishable from selecting deck *B* in long-term average expectations. Similarly, selecting deck *C* is indistinguishable from selecting *D*. For the conflict version, the feasible region and Pareto front are identical which means that every possible decision-maker is Pareto optimal. Moreover, the feasible regions for our three decision-makers (i.e. soft-max, epsilon-greedy, and selective attention) are identical and therefore are indistinguishable in long-term average expectations. For the non-conflict version, there exists an action (deck *C* or *D*) that optimizes both of the objectives simultaneously.

**Fig. 2.**
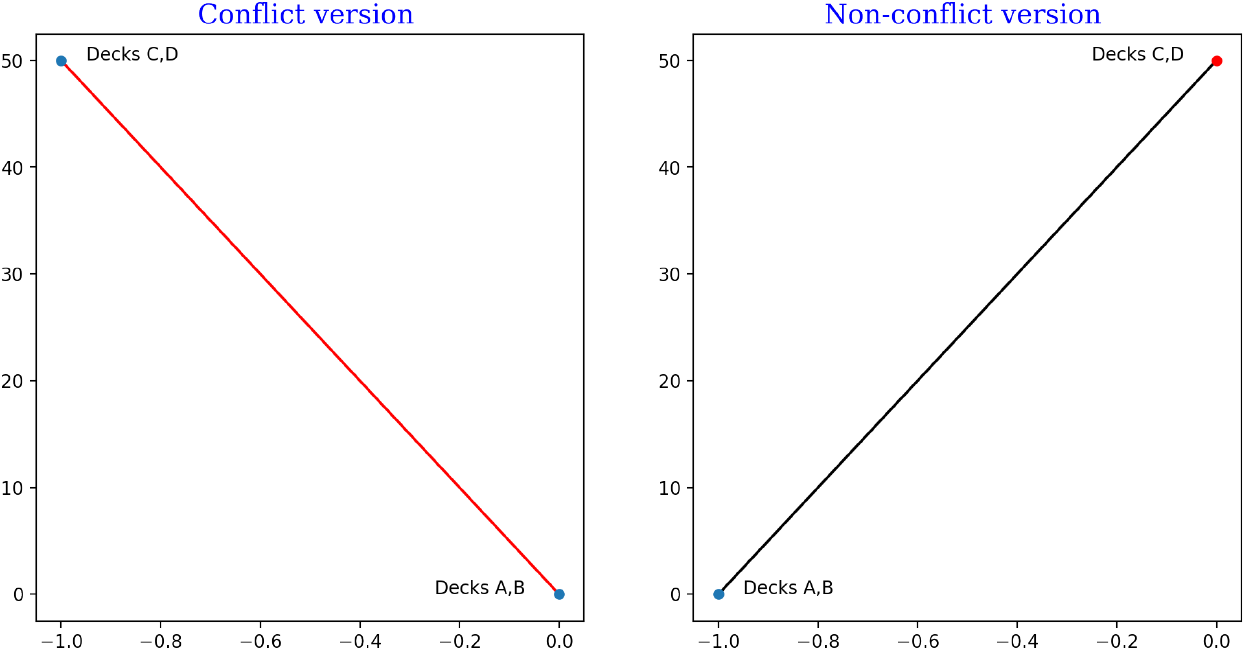
Feasible regions of all decision-makers for a conflict and non-conflict version of the spider gambling task. Pareto Front is in red and feasible region is in black. In the conflict version, the Pareto front and the feasible region coincide, meaning that all decision-makers are Pareto optimal.

### 4.2 Weather task

In Aupperle et al. (2011), the authors designed an AAC task to study how self-reported anxiety is related to approach behavior and how it differed between males and females. The task consisted of four phases with increasing level of conflict with the initial phase having zero conflict. When a participant selects an action, it could either result in a numerical reward and a negative affective stimuli (a disturbing image) or no numerical reward and a positive affective stimuli (pleasant image), thus creating a conflict except in the initial phase when the numerical reward was zero (avoidance only). As shown in Figure 3, the feasible region of all decision-makers collapses to a single line segment and coincides with the Pareto Front, except in the initial phase. This means that all decision-makers in conflict phases in this task will be Pareto optimal. Further, the feasible regions for our three decision-makers are again identical.

**Fig. 3.**
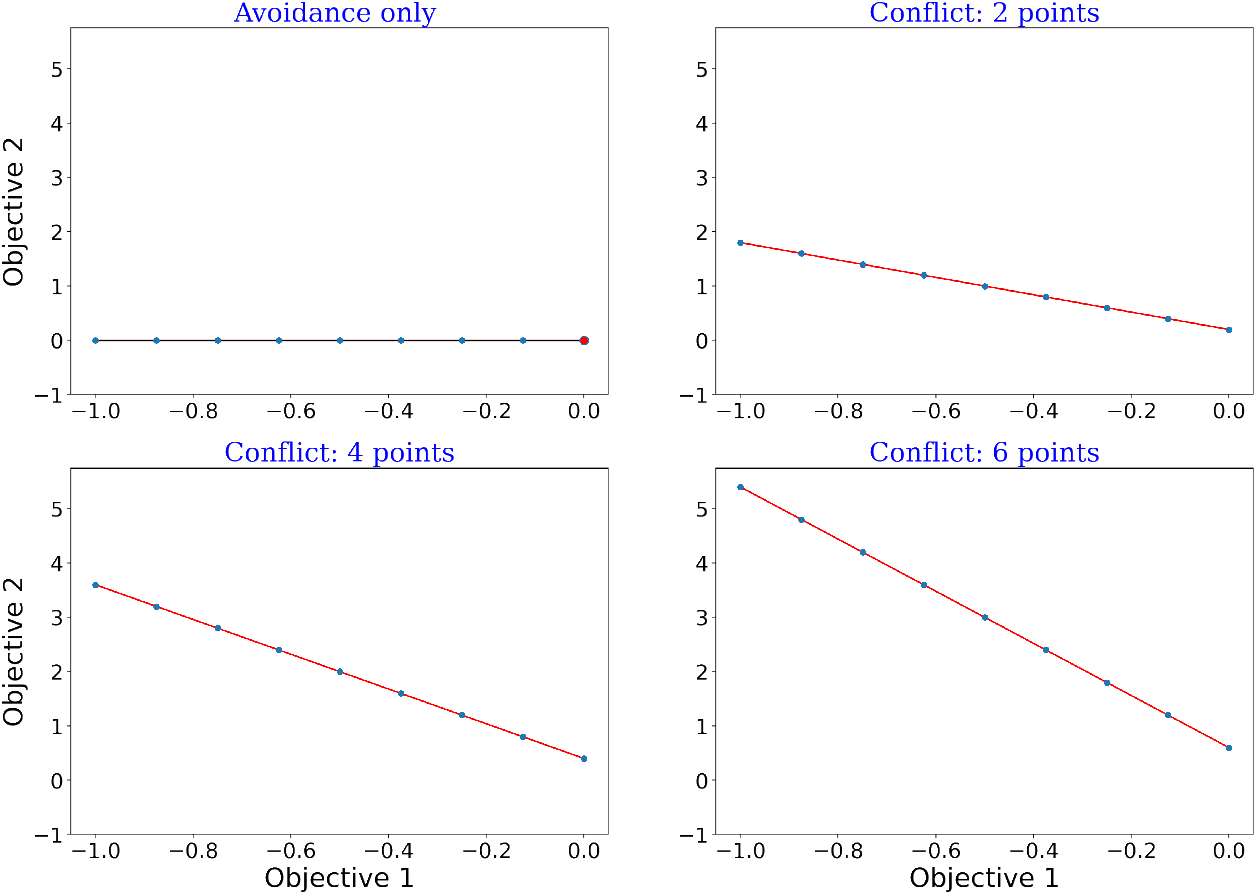
Feasible regions of all decision-makers for four phases of the weather task (Aupperle et al., 2011). Pareto Front is in red and feasible region is in black. Except for the initial phase (avoidance only), Pareto Front coincides with the feasible region, meaning that all decision-makers are Pareto optimal.

### 4.3 Trauma-related AAC task

In Weaver et al. (2020), the authors introduced the trauma-related AAC task in order to investigate how individuals with post-traumatic stress disorder differ from controls in their decision-making during AAC. The trials belonged to either a *conflict* phase where the action most likely to win points was most likely to show a trauma-related image or to a *congruent* where the action most likely to win points was least likely to show a trauma-related image.

The reward structure Ω of the trauma-related AAC task was separated into four phases (two conflict phases and two congruent phases) with 75 trials of each type. At a trial in a given phase, the participants choose from three shapes (circle, square and hexagon), each having a phase-specific chance of showing a trauma-related image and positive point reward. For instance, if the hexagon is selected during block 1 of the conflict phase, it would have an 80% probability of showing a trauma-related image and 80% probability of yielding a positive point reward. On the other hand, the same hexagon would have 20% probability of presenting trauma-related image and 80% chance of positive point reward. In Figure 4, we notice again that for the conflict versions of the AAC task, the Pareto front equalled the feasible region 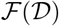 of all decision-makers, which means that all decision-makers in this setting would be Pareto optimal.

**Fig. 4.**
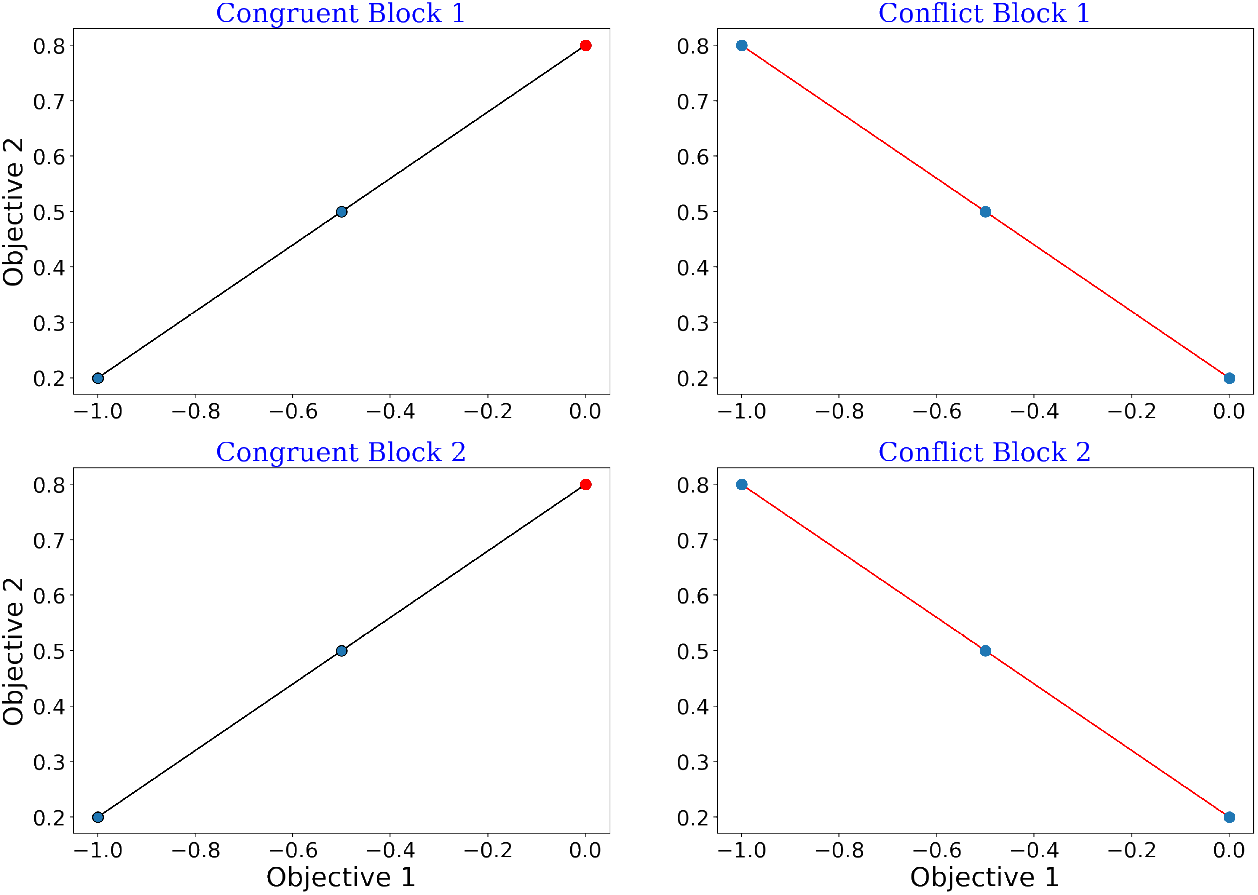
Feasible regions of trauma-related AAC (trAAC) task. In the congruent blocks, Pareto Front (red) is a proper subset of the feasible region (black), indicating some actions dominate others. In contrast, Pareto Front (red) coincides with the feasible region for the conflict phases, meaning all actions are Pareto Optimal.

To conclude this section, we note that the conflict phases or versions of AAC tasks we considered from psychology literature all had the property that the Pareto front precisely equalled the feasible region of all decision-makers. This means that all decision-makers one could consider would be Pareto optimal. In the next section, we will show that once we consider an AAC task where the Pareto front is a proper subset of the feasible region, a range of interesting behaviors can be observed for different decision-makers.

## 5 Potential AAC tasks

If we consider an AAC task with a feasible region that is not one-dimensional, then more interesting phenomena arises. Figure 5 shows a potential AAC task with 4 actions, each of which are Pareto optimal. The Pareto front (shown in red) lies on the boundary of the feasible region 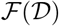 of all decision-makers, but is not equal to this feasible region. Therefore, there will be decision-makers who behave sub-optimally. Further, the feasible regions of different type of learners differ from each other, which was lacking for the AAC tasks we analyzed from the literature. Since all 4 vertices are Pareto optimal, the feasible region of the epsilon-greedy decision-maker is the union of the seven line segments connecting the centroid of the vertices of 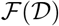 to either a vertex or a midpoint on the Pareto front between two vertices.

**Fig. 5.**
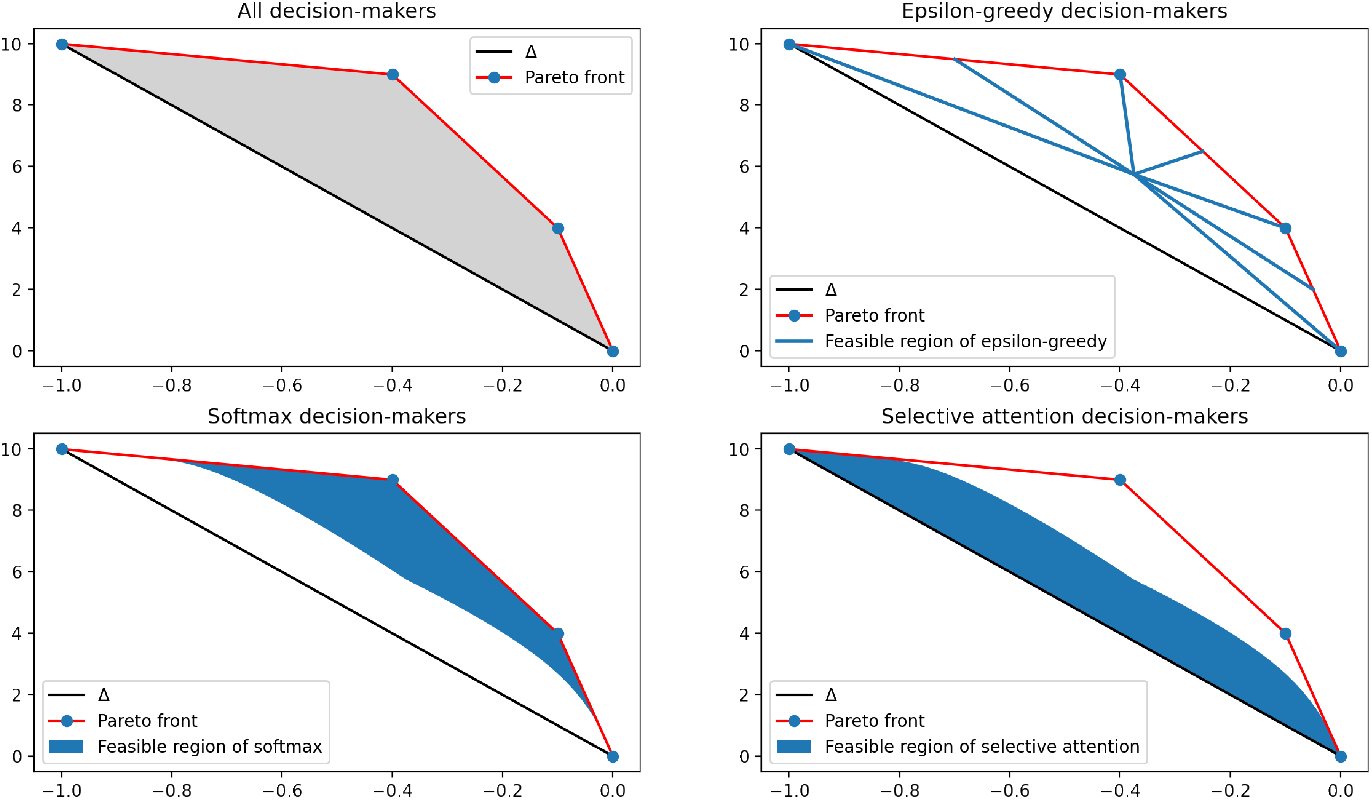
Example of an AAC task that has four vertices of the feasible region 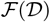 on the Pareto front.

In comparison, the softmax and selective attention decision-makers have feasible regions that are not a finite union of line segments. The feasible region of the softmax decision-maker has the Pareto front on its boundary. The part of the boundary that is not the Pareto front is exactly the curves obtained by setting either *w*_1_ = 0 or *w*_2_ = 0. For example, the curve for *w*_1_ =0 captures decision-makers who ignore the first reward dimension and connects the vertex corresponding to always selecting the action that maximizes the second reward dimension to the centroid of vertices of 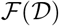.

Meanwhile, these curves for *w*_1_ = 0 or *w*_2_ =0 are also part of the boundary of the feasible region of the selective attention decision-maker, capturing selective attention decision-makers whose respective parameters are also set to *w*_1_ = 0 or *w*_2_ = 0. This boundary, however, is the only thing shared by the two feasible regions. The feasible region of selective attention decision-makers contains the remaining part of 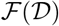 not in the feasible region softmax decision-maker. Unlike softmax decision-makers which are drawn toward the Pareto front with increasing τ, selective attention decision-makers are drawn toward the line segment Δ connecting the vertices of 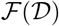 that maximize one of the two objectives.

While the previous example shows that softmax and selective attention decision-maker can behave quite differently (as indicted by the non-overlapping feasible regions except at their boundaries), this is not the case in general. Consider an example of an AAC task where, instead of the Pareto front being different from Δ, the Pareto front and Δ are the same. Then, we should expect the feasible regions of the two decision-makers to be identical. Figures 6 and 7 confirms this expectation.

**Fig. 6.**
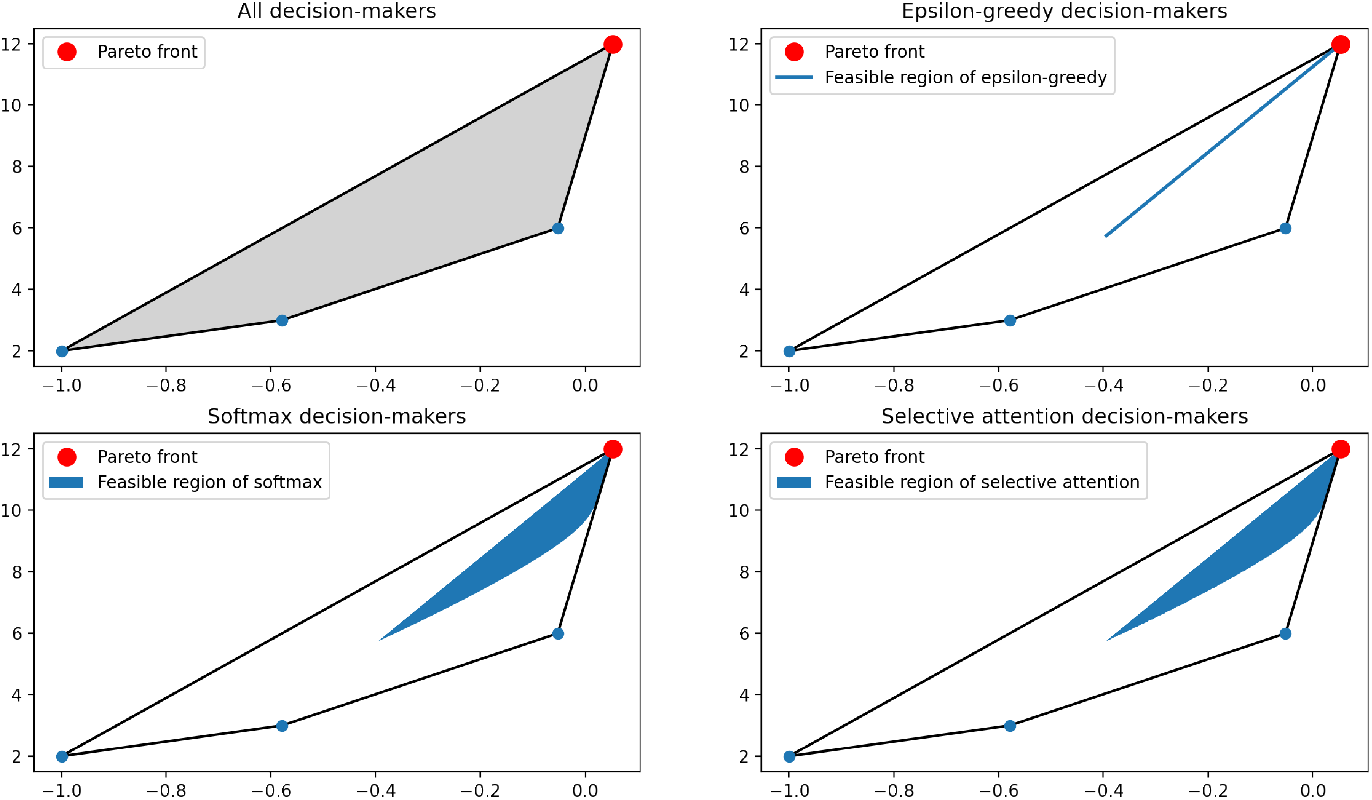
Example of an AAC task that has one vertex of the feasible region 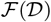 on the Pareto front

**Fig. 7.**
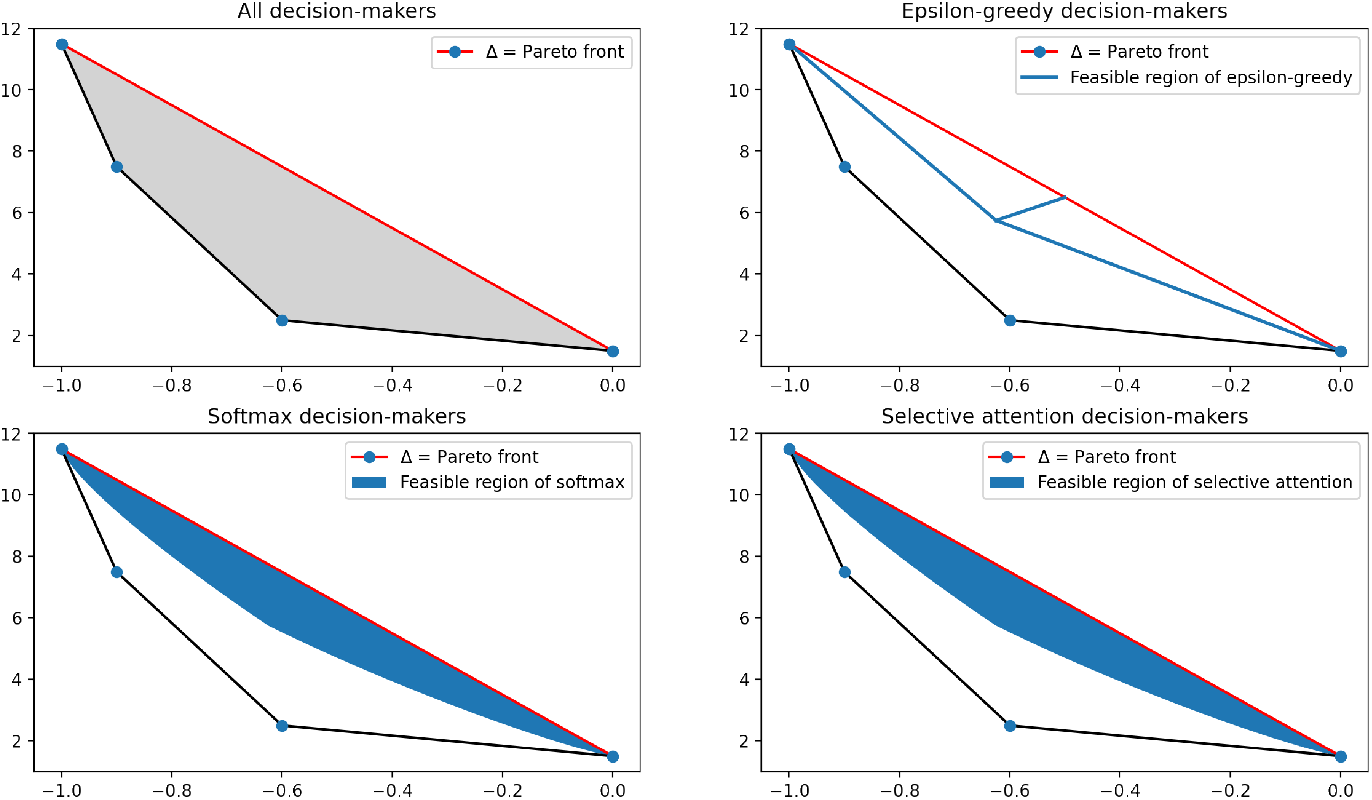
Example of an AAC task that has two vertices of the feasible region 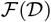 on the Pareto front

Notice that the Pareto front always contains the vertices that optimize each objective. Therefore, the above scenario occurs if and only if every Pareto optimal action maximizes at least one of m objectives. This generalizes an observation from the section on prior AAC tasks (Section 4): that even for an AAC task with a two-dimensional feasible region 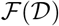 of all decision-makers, the families of decision-makers we consider can still be indistinguishable when the Pareto front is the same as Δ. Lastly, Figure 8 shows a final example of an AAC task, in which the interior of the feasible regions of softmax and selective attention decision-makers can overlap without being identical.

**Fig. 8.**
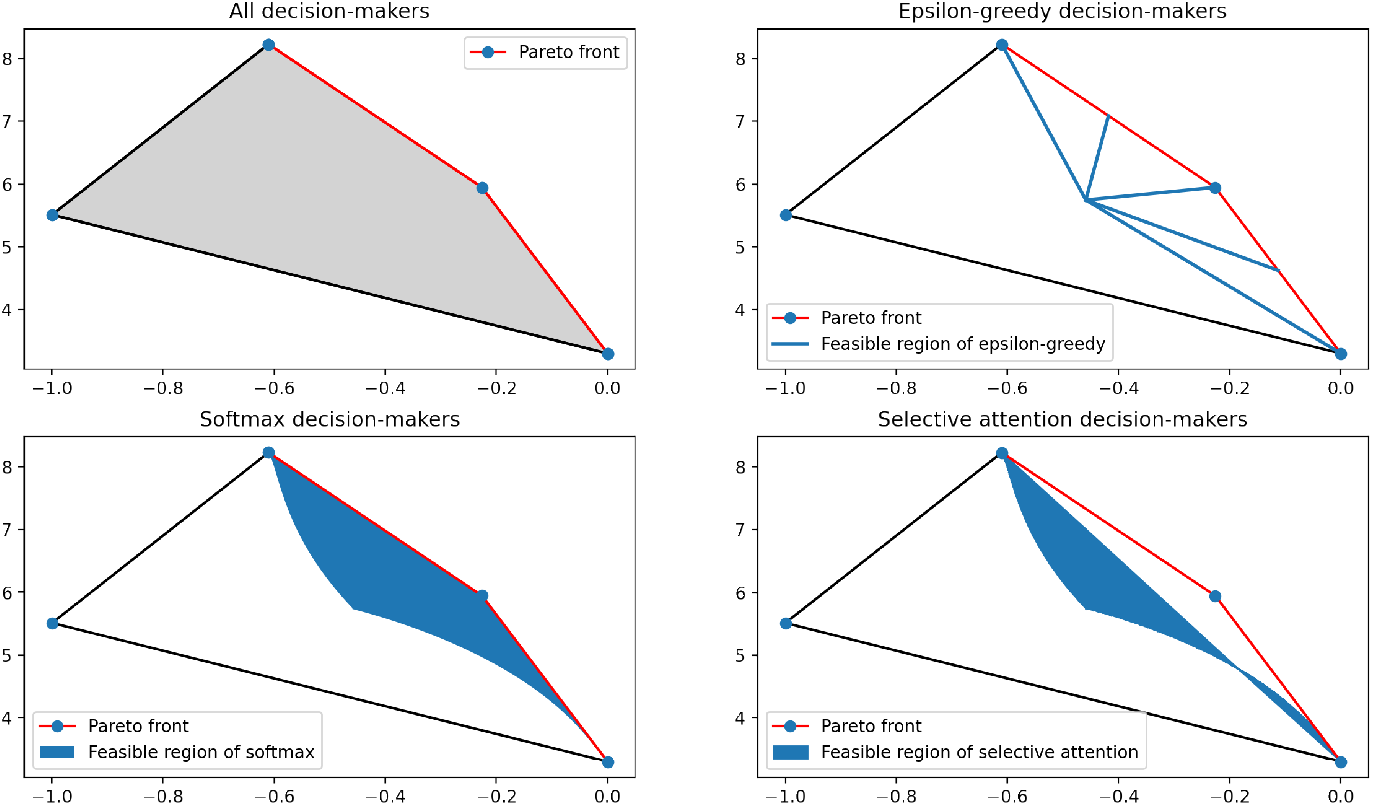
Example of an AAC task that has three vertices of the feasible region 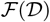 on the Pareto front

Before we conclude this section, let us mention another potential way to look at and compare these decision-makers. So far, the way we have been comparing decision-makers is to fix the reward structure Ω of an AAC task and then look at feasible regions of decision-makers which is obtained by considering all possible values of their parameters (i.e., *τ*, ***w*** for the softmax decision-maker.) While this analysis gives us some insights into differences between these decision-makers, it says nothing about particular instances of them for given parameters. More specifically, given an AAC task and an instance of a decision-maker, captured by a point in the feasible region 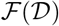, how can one tell if, for example, it is a softmax decision-maker or an epsilon-greedy one?

To this end, we produced Figure 9 by perturbing each vertex ***μ**_i_* of the feasible region 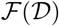 and plotting the change in long-term average expected rewards of three specific decision-makers (one from each family). For instance, one can see from the top-right panel that the decision-makers were much more sensitive when a vertex that’s closest to the fixed decision-maker is perturbed. Therefore, it may be advisable to perturb other vertices to get distinct behaviors from the different families of decision-makers.

**Fig. 9.**
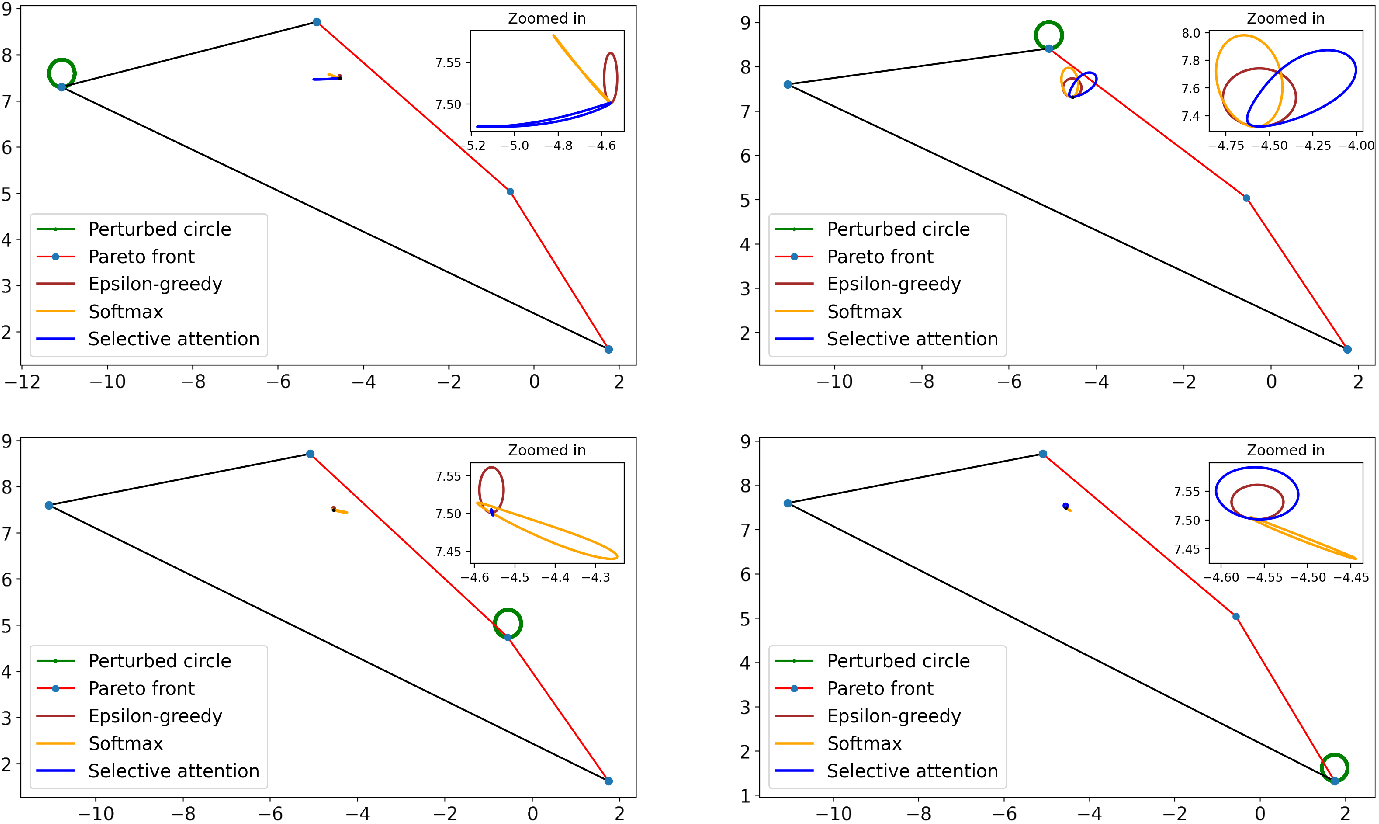
Sensitivity of decision-makers to perturbation of the reward structure for an AAC.

In the bottom-left panel, the selective attention decision-maker has barely any change compared to the other two decision-makers. This behavior is expected from our previous observations, since, like the AAC task in Figure 8, the feasible region of the selective attention decision-maker did not include the vertex being perturbed. Put differently, since the perturbation did not affect the line segment Δ, the selective attention decision-maker remained largely unaffected by this perturbation.

On the other hand, if we look at the top-left panel,the selective-attention decision-maker can still be relatively sensitive when a vertex ***υ*** not on Δ is perturbed. However, this maybe due to the fact that the said perturbation causes **υ** to become a close contender to be on Δ.

## 6 Discussion

In this paper, we investigated approach-avoidance conflict (AAC), an important phenomena described in the psychiatry and psychology literature, from the perspective of multi-objective, multi-armed bandits (MO-MAB) (Drugan & Nowe, 2013). The crux of AAC is that making a decision, such as whether to approach or avoid, is fundamentally multi-objective: the decision that maximizes desirable outcomes (reward) might not be the decision that minimizes undesirable outcomes (harm). In this vein, the MO-MAB provided a suitable framework to study AAC, since it described a decision-making scenario that is multi-objective. Specifically, current decision-making tasks elicit an AAC by pairing *n* different options for the participant to choose from on a given trial with *m* different outcomes (some rewarding and some harmful). We modeled such an AAC task by a MO-MAB with *n* actions, each bringing an independent outcome vector of *m* dimensions.

With this setup, we introduced the definition of a general decision-maker as a limiting sequence of decisions during an AAC task. This definition allowed us to disentangle the decision process from the learning process so to focus on the former. Importantly, we could analyze decision-makers by studying their longterm average expected outcomes, which is a point in 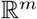. Therefore, comparing two different decision-makers amounts to comparing two points in 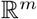, which could then be naturally compared using the concept of Pareto dominance. This motivated us to define *feasible region* for a collection of decision-makers to be the corresponding set of long-term average expected outcomes. We showed that the feasible region of all decision-makers is a polytope defined by the *n* expected outcome vectors in 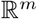associated with the *n* actions and that the Pareto front of this feasible region plays an important role in distinguishing between different types of decision-makers.

We also introduced three types of decision-makers with the intention that researchers could use these models to explain decision-making data in future AAC experiments. These examples extend a TD learning model of human decision-making, already ubiquitous in single-objective settings, to the multi-objective setting. In general, a TD learning model combines a learning process such as *Q*-learning and a decision-making process such as epsilon-greedy. When there are m different objectives, we suggest that a decision-maker could learn *m* different action-value estimates or *Q* tables (one for each outcome dimension) and then use *scalarization*, a common method in the MO-MAB literature (Drugan & Nowe, 2013), to combine *Q* values across all the outcome dimensions in some way to obtain a single estimate. We specifically propose decision-makers that use linear scalarization to turn the *m* different *Q* tables to a single estimate through a weighted sum with a non-negative, normalized weight vector ***w*** = (*w*_1_, *w*_2_,…,*w_m_*)^*T*^ and then pass the weighted sum into one of two popular decision-making rules: softmax or epsilon-greedy. We also proposed a third example of a decision-maker that does not use scalarization; instead, it presumes that a decision-maker selectively attends to one of the reward dimensions at a time with a certain probability and applies a softmax rule to the *Q* values of the attended dimension. We can then show that under suitable updates of the *Q* values, we can identify a decision-maker of any of the three types, substantiated by their parameters, with the point in 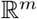 representing their long-term average expected outcomes.

We applied our framework to model three previous AAC tasks in Aupperle et al. (2011); Pittig et al. (2014); Weaver et al. (2020). One thing became clear: the Pareto front equals the whole feasible region during the phases or versions of the task with an AAC. The main reason why this happened is that the feasible regions of all decision-makers has zero volume in 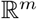 for these AAC tasks. Specifically, the feasible regions were line segments in 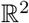 and so were the Pareto fronts. As a result, one is unable to distinguish between the feasible regions of the three examples of decision-makers (i.e., softmax, greedy epsilon, and selective attention). Further, all decision-makers are Pareto optimal, e.g., an individual who avoids the spider decks at all costs is equally optimal as someone who maximizes their monetary rewards (Pittig et al., 2014). If one ascribes to the viewpoint that psychopathology arises from computational deficits (see Haynos, Widge, Anderson, and Redish (2022) for an overview), then these previous tasks may be better suited for identifying other deficits (e.g., overvaluation of harm) than the specific deficit of making sub-optimal decisions during an AAC.

To resolve this issue, we introduced a set of AAC tasks where the Pareto front is a proper subset of the whole feasible region and showed that the three examples of decision-makers (softmax, selective attention and epsilon-greedy) can have distinctive feasible regions. These tasks are designed by specifying mean outcomes that yield a feasible region of all decision-makers that has positive volume in 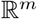. Once we have that, we saw that the three different decision-makers can have feasible regions that are either identical, non-overlapping except at the boundary, or something else entirely. Propositions 2, 3 and 4 allow us to predict each of these cases as a function of the Pareto front and the hyperplane Δ connecting the vertices in 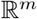 corresponding to actions that maximize mean outcomes along at least one of the outcome dimensions.

If it is desirable to distinguish between a softmax and selective-action, our findings support an AAC task which includes Pareto optimal actions that do not maximize mean outcomes along at least one of the outcome dimensions. Further, if it is desirable to distinguish between single instances of decision-makers, we show how one could perturb a vertex along the Pareto front, especially a Pareto optimal vertex that does not maximize any of the objectives.

There are several limitations of this paper that one should consider. One limitation is that there are alternative models that have been proposed in the literature to describe human decision-making during an AAC. Smith et al. (2021a, 2021b), for example, use partially observable Markov decision processes to model the weather AAC task of Aupperle et al. (2011), which, while arguably more difficult to implement that the models we considered, may turn out to be a better fit to data. Meanwhile, Zorowitz et al. (2020) used the beta-pessimistic *Q*-learning of Gaskett (2003) to capture AAC elicited during the balloon analogue risk task of Lejuez et al. (2002). Though it is important to note that, since this task elicits an AAC through the *chance* of either a desirable or undesirable outcome as opposed to the *simultaneous occurrence* of both types of outcomes, this model would have to be adapted to handle the multi-dimensional outcomes of the AACs considered in the present paper. Additionally, there are other AAC tasks in the literature that we could have considered, such as the aforementioned balloon analogue risk task of Lejuez et al. (2002) or the predator-forager task of Bach et al. (2014).

Another limitation is the collection of assumptions underlying our model. First, we treat outcomes of each objective as a numerical value, regardless of whether they are sound cues, disturbing images or electric shocks. Second, we identified a decision-maker with their long-term average expected rewards. Alternatives are to identify a decision-maker with either their long-term discounted reward or their decisions in the short-term. Long-term discounted rewards, for example, could capture a range of behaviors based on how much the decision-maker values short-term versus long-term rewards. Third, we focused on using decision-making rules that have a precedence in the psychology literature (i.e., epsilon-greedy, softmax and selective attention) and used Pareto optimality to compare decision-makers. One can instead use quality indicators such as the hypervolume metric of Zitzler, Knowles, and Thiele (2008) to compare decision-makers or use Chebyshev scalarization of Van Moffaert and Nowé (2014) instead of linear scalarization to obtain the scalarized *Q*-table. A final limitation is that, in an effort to provide simple insights for future study, we focused entirely on the decision-making part of human decision-making during a bandit task. Thus, our framework may miss interesting decision-making behavior that arises while learning during AAC or while using a non-bandit task for which current decisions impact future contingencies.

In conclusion, we introduced a mathematical framework to analyze, model data from, and design AAC tasks. We also derived properties of these models for describing human decision-making during AAC. This framework provides a pathway forward for researchers looking to test for sub-optimal decision-making or to distinguish between approaches to decision-making among participants in an AAC task.

## 7 Statements and Declarations

### 7.2 Data availability

Data sharing not applicable to this article as no datasets were generated or analysed during the current study. Code is publicly available at https://github.com/eza0107/Multi-Objective-RL-and-Human-Decision-Making.

## Appendix A Proof of long-term limit

We begin by proving the claim made in Definition 1. That is, for a decision-maker *D* associated with action sequence 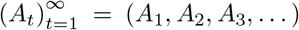, one has:

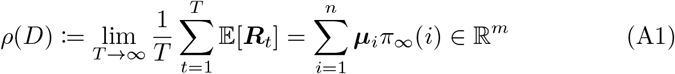

*Proof* Let *π_t_* and *π*_∞_ be the probability distributions of *A_t_* and *A*_∞_ as defined in Definition 1. Then,

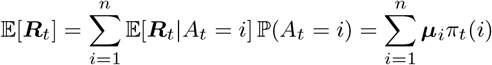

Note that *n* is fixed and finite and that *π_t_*(*i*) → *π*_∞_(*i*) for each 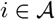, as *t* → ∞. Therefore,

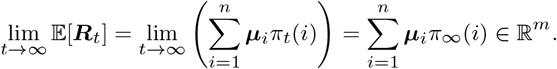

Now the conclusion directly follows from the Stolz-Cesaro theorem of Stolz (1885):

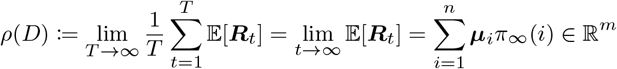

## Appendix B Proof of Proposition 1

Here, we give the proof of Proposition 1, which says that

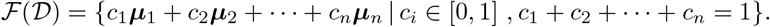

*Proof* The proof in Appendix A tells us that for a given decision-maker *D*, *ρ*(*D*) ∈ {*c*_1_***μ***_1_ + *c*_2_***μ***_2_ + … + *c_n_**μ**_n_* | *c_i_* ∈ [0, 1], *c*_1_ + *c*_2_ + … + *c_n_* = 1} by simply choosing *c_i_* = *π*_∞_(*i*). Therefore,

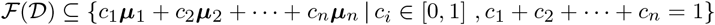

For the other direction, any given 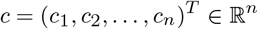 with *c*_1_ + *c*_2_ + … + *c_n_* = 1, *c_i_* ≥ 0 defines a probability distribution over the action set 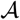. If we let *D* be a decision-maker who chooses action *i* with probability *c_i_* on every trial, then we get:

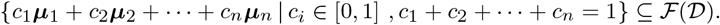

## Appendix C Proofs of Propositions 2–4

Before we move onto the proofs of the main propositions below, let’s briefly elaborate on the Pareto front 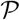. As we defined in the main text, 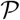 is precisely the Pareto optimal part of the boundary of the convex polytope 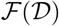. In other words, 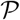 is the finite union of the Pareto optimal faces of 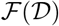, each of which is a convex polytope of at most *m* − 1 dimension. If *F* is one such face of dimension, then by renumbering vertices we can represent

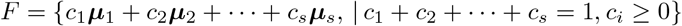

with *s* ≤ *m*. Importantly, ***μ***_1_,…,***μ**_s_* will be linearly independent from our assumed non-degeneracy of Ω (Assumption 1). Furthermore, let us note that no vertex ***μ*** on 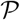 is dominated by any convex combination of the vertices in Ω. That is, if we let 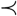 denote Pareto dominance, then

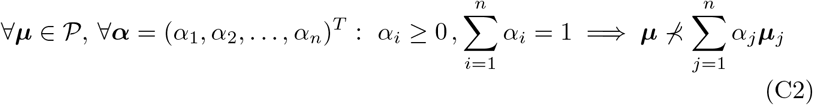

But if we restrict our attention to F only, something even stronger is true:

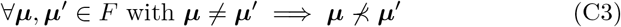

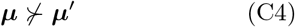

Now we prove an auxiliary lemma that is a strengthening of the above property for a given face 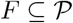.

**Lemma 1** Let *F* be a face on 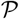, generated by convex combinations of linearly independent, Pareto optimal vertices ***μ***_1_,…,***μ**_s_*. Then for any *i* ∈ {1, 2,…,*s*}, one has:

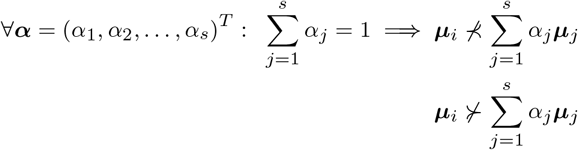

*Proof* Note that the strengthening refers to the removal of the requirement that *α_j_*’s must be non-negative. This means that none of ***μ**_i_* is dominated by an *affine* combination of ***μ***_1_,…,***μ**_s_*. Suppose otherwise and assume without loss of generality that ***μ***_1_ is strictly dominated by an affine but not convex combination of the ***μ***_1_,…,***μ**_s_*:

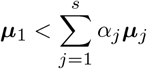

and vice versa. Since at least one of the *α_j_* must be negative and another of the *α_j_* must be positive, we can alternatively write the above as:

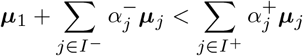

with

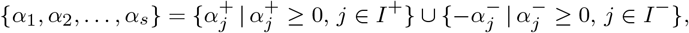

and *I*^+^ ∪ *I*^−^ = {1,2,…,*s*}, *I*^+^ ∩ *I*^−^ = *ϕ*. Notice that:

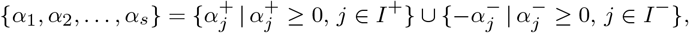

Therefore, we obtain:

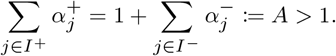

However, this leads to a contradiction to (C2) since the convex combination of ***μ***_1_ and {***μ**_j_*: *j* ∈ *I*^−^}, a Pareto optimal point on the face *F*, cannot be dominated by a convex combination of {***μ**_j_*: *j* ∈ *I*^+^}. The inequality in the other direction follows the exact same reasoning.

We need another auxiliary result, namely the famous Farkas’ lemma which we state here without proof.

**Lemma 2** (Farkas’ lemma) Let 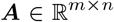 and 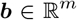. Then exactly one of the following two assertions are true:

1. There exists an 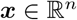 such that ***Ax*** = ***b*** and ***x*** ≥ 0.
2. There exists an 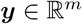 such that ***A**^T^****y*** ≥ 0 and ***b**^T^****y*** < 0.

Now we are ready to state and prove our final auxiliary lemma:

**Lemma 3** Let *F* be a Pareto optimal face with vertices ***μ***_1_,…,***μ**_s_*. There exist strictly positive 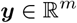 such that

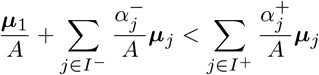

and for *j* = *s* + 1,…, *n*,

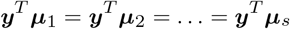

*Proof* From Proposition 1, the feasible region 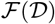 is a compact, convex polytope. As such, it can be expressed as the intersection of a finite number of half-spaces (Weyl-Minkowski theorem):

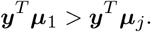

Consider indices of those half-spaces for which ***μ***_1_,…,***μ**_s_* are on its boundary:

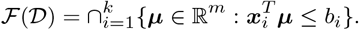

This set, in fact, characterizes the face *F*:

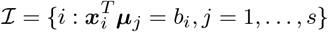

Use 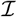 to define the closed, non-empty, convex set:

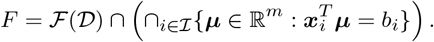

Convexity follows since *C* is the intersection of a finite set of half-planes. Critically, *C* does not contain any non-negative vectors, except for **0**; if it did, we could move ***μ***_1_ in the direction of this non-negative, non-zero ***ν*** ∈ *C* to get a point 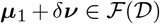 for small δ > 0 that Pareto dominates ***μ***_1_, contradicting ***μ***_1_ being a vertex of a Pareto optimal face. Consequently, we can find a hyperplane, with normal ***z***, to separate *C* from the (compact, non-empty, convex) set of convex combinations of standard basis vectors *e_i_* in 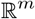:

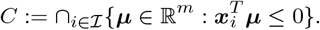

The last inequality says each entry in ***z*** is strictly positive, and hence, ***z*** is strictly positive.

We will use the fact that the vector ***z*** can be expressed as a conic combination of the vectors ***x**_i_* for 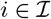:

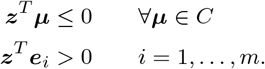

This follows from Farkas’ lemma, since otherwise we could find 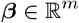 such that

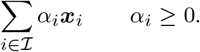

The first inequality says that ***β*** lives in *C*, but then the second equality contradicts the condition that ***z***^*T*^***μ*** ≤ 0 for all ***μ*** ∈ *C*.

We are now ready to construct ***y***. Let

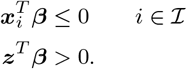

for *∊* > 0 sufficiently small so that ***y*** is strictly positive. Such an e exists, since ***z*** is strictly positive. Further, our characterization of ***z*** above allows us to express **y** in the form 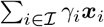 for some strictly positive *γ_i_*. Therefore,

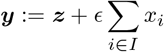

since for each *j* = 1,…, *s*,

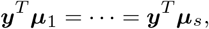

It is also the case that, for each *j* = *s* + 1,…, *n*,

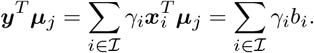

since

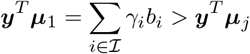

for some 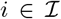. Otherwise, ***μ**_j_* would also be on *F*, which contradicts the initial assumption that ***μ***_1_,…,***μ**_s_* are all the vertices on *F*. We can conclude that **y** has the desired properties.

Now we are ready to prove the propositions:

*Proof of Proposition 2* Fix the parameters *ε* and ***w***. Recall that epsilon-greedy decision-maker samples uniformly at random with probability *ε* from all the vertices and with probability 1 − *ε* choose a vertex that maximizes ***w**^T^**μ**_i_* from 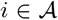. Therefore, we can write the probability of action *i* being chosen as:

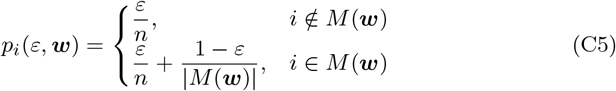

where

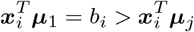

Our epsilon-greedy decision-maker with the given parameters is characterized by:

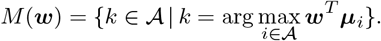

which trace out the line segment connecting 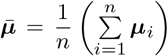, the centroid of the vertices of Ω, and 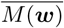, the centroid of the vertices with indices in *M*(***w***).

First note that *M*(***w***) must be the indices of vertices of some Pareto optimal face *F*. To see this, suppose ***μ***_1_, ***μ***_2_,…***μ**_s_* are the vertices with indices in *M*(***w***), but not the vertices of any Pareto optimal face. Then by nature of not being the vertices of a Pareto optimal face, there must exist a convex combination ***μ*** of these vertices that is *strictly* dominated by a point ***μ***″ on the Pareto front:

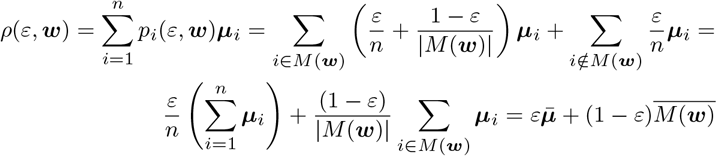

Together with ***w*** ≥ 0, ***w*** ≠ 0, ***w**^T^**μ***_1_ = … = ***w**^T^**μ**_s_*, and ***μ*** being a convex combination of ***μ***_1_,…, ***μ**_s_*, this implies that

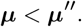

Meanwhile, ***μ*″** would be itself a convex combination of vertices ***μ***_*n*1_,…, ***μ***_*n*__*k*_ on the Pareto front:

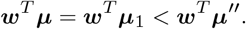

and so,

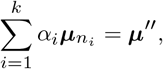

Since *α_i_* ≥ 0, it follows that for at least one index *i*, one has ***w**^T^**μ***_*n*__*i*_ > ***w**^T^****μ***_1_, which contradicts 1 ∈ *M*(***w***). What this shows is that *M*(***w***) must be the indices of vertices of some Pareto optimal face *F*.

Now let *F* be a Pareto optimal face of 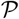 with vertices ***μ***_1_,…,***μ**_s_*. Our goal is to construct ***w*** so that the set *M*(***w***) is exactly the indices {1,…,*s*}. From Lemma 3, we can find strictly positive ***y*** such that

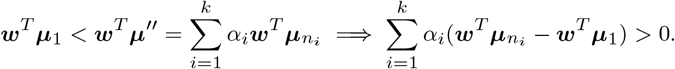

and for each *j* = *s* + 1,…,*n*,

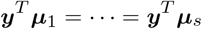

Rescaling ***y*** by a positive number does not change the equalities and inequalities above. Therefore, letting ***w*** = ***y***/∥***y***∥_1_ gives us *M*(***w***) = {1,…,*s*}, as desired.

We now have that there is some **w** ≥ 0 with ∥***w***∥_1_ = 1 such that *M*(***w***) = {1,…,*s*} if and only if ***μ***_1_,…,***μ**_s_* are vertices of a Pareto optimal face. This completes the proof, since now we know that the feasible region of epsilon-greedy decision makers is characterized by line segments between the centroid of the vertices of the feasible region and the centroid of the vertices of each Pareto optimal face. Note also that any Pareto optimal vertex is itself a Pareto optimal face due to Assumption 1.

*Proof of Proposition 3* Consider the space

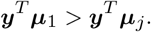

Parameters *τ*, ***w*** for a softmax decision-maker can then be specified by selecting any (*τ*, ***u***) from 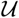 and then letting w have ***u*** as its first *m* − 1 entries and 1 − ∥***u***∥_1_ as its remaining entry. The proof will be divided into two parts. In the first part, we show that if Ω is not degenerate and has a non-empty interior, then the softmax decision-maker *ρ*(*τ*, ***w***) maps an interior point of 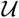 to an interior point of Ω. This will mean that any boundary points of the feasible region for softmax decision-makers will be generated from either the boundary of 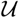 (i.e. when one of the w-are zero as stated in the proposition) or from taking the limit as *τ* → ∞. In the second part, we will prove that any point on the Pareto front can be approximated by a sequence of interior points of 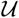 in the limit *τ* → ∞.

For the first part, we assume that Ω has a non-empty interior. Fix 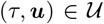, define ***w*** accordingly, and do a linear change of variables *τ****w*** = ***x*** (only for this part of the proof):

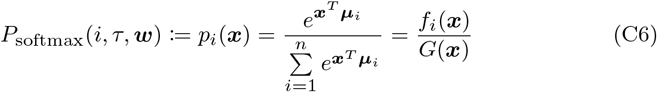

where 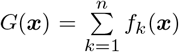. Note that this change of variables, from the interior of 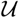 to (0, ∞)^*m*^, is a local homeomorphism, since it has a non-vanishing Jacobian:

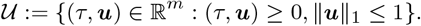

With this setup, consider the mapping from ***x*** to the long-term average expected rewards for the decision-maker with parameter ***x***:

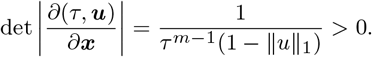

Now for the actual proof, we first show that the Jacobian of ***F*** is positive semi-definite for ***x*** > 0. In fact,

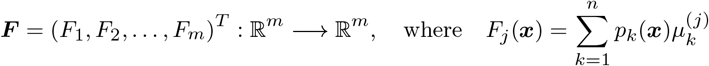

and therefore:

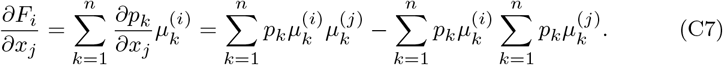

For given ***x*** > 0, *p*_1_(***x***), …, *p_n_*(***x***) defines a discrete probability distribution on the vertices of Ω. Namely, there exists a random vector ***X*** = (*X*_1_, *X*_2_,…, *X_m_*)^T^ which takes the value ***μ**_i_* with probability *p_i_*(***x***) for *i* = 1, 2,…,*n*. That is:

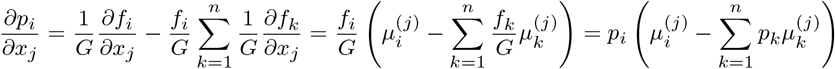

The component *X_i_* then takes the value 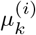 with probability *p_k_*, and therefore:

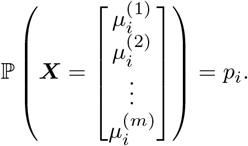

Furthermore,

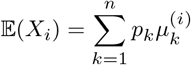

Putting everything together, we obtain:

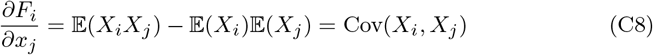

This means that the Jacobian matrix is:

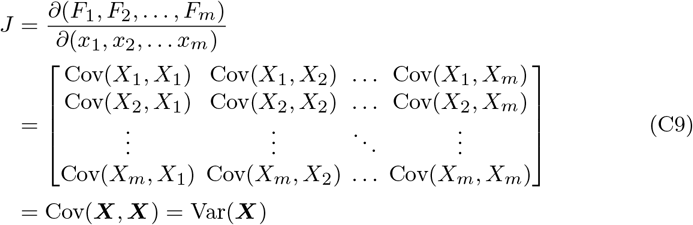

It’s well-known that the covariance matrix is positive semi-definite but we claim that this is positive definite. If it were not positive-definite, then there exists a nonzero vector 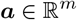 such that Var(***X***)***a*** = 0. Therefore,

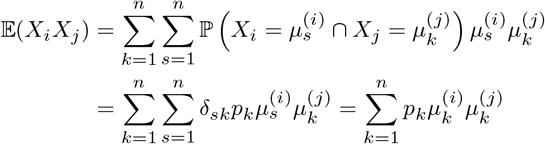

This means with probability 1, the vector ***X*** lives on some hyperplane ***a**^T^**X*** = *b*, which would be impossible unless Ω has zero volume in 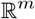, contradicting our assumption that Ω has non-zero volume. To see this, a simple computation of the variance yields:

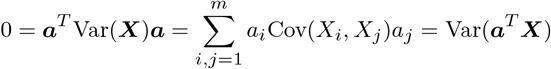

Since *p_i_* > 0, this means that:

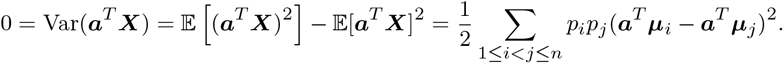

Therefore, the Jacobian is non-vanishing on the interior of the domain [0, ∞)^*m*^, and our map ***F*** is locally a homeomorphism on [0, ∞)^*m*^. In addition, we have already shown changing variables from (*τ*, ***u***) in the interior of 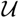 to (0, ∞)^*m*^ is a local homeomorphism. Therefore, the composition of these two local homeomorphisms is a local homeomorphism. We can conclude that *ρ*(*τ*, ***w***) maps interior points of 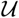 to interior points of Ω.

For the second part of the proof, we pick a point on ***μ*** on the Pareto front. It will belong to a Pareto optimal face, *F*, which is generated by a finite number of vertices 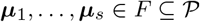. That is, we have:

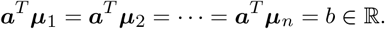

We can assume that the vectors ***μ***_1_,…,***μ**_s_* ∈ *F* are linearly independent (Assumption 1) and the maximal set of vectors that generate *F*. Recall that:

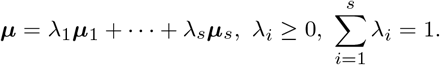

We will show that a limiting sequence of (*τ*, ***w***) have *P_i_*(*τ*, ***w***) limiting to λ_*i*_ for *i* = 1,…,*s* and to zero for *i* = *s* + 1,…,*n*. The corresponding sequence of points in Ω will have

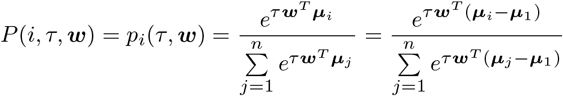

which would complete the second part of our proof.

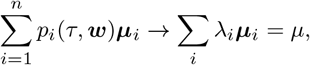

Without loss of generality, assume λ_1_ ≥ λ_2_ ≥ … ≥ λ_*k*_ > 0 and λ_*i*_ = 0 for *k* < *i* ≤ *s*. Consider *α* > 0. If *s* = 1, let ***z*** be any non-negative, non-zero vector. If *s* > 1, let us find non-negative, non-zero 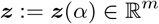 that solves

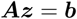

where

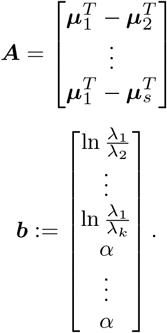

To show that such a solution ***z*** exists, we will prove that the second assertion in Farkas’ lemma is violated. Let us pick some 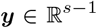 such that ***b**^T^**y*** < 0. This implies **y** ≠ **0** and if we let ***y**^T^* = (*y*_2_, *y*_3_,…***y**_s_*), then:

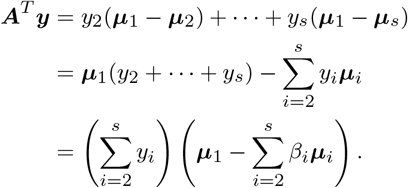

If *y*_2_ + *y*_3_ + … + *y_s_* ≠ 0, then let

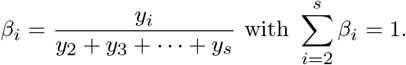

We know from Lemma 1 that neither of ***μ***_1_ and 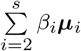 can Pareto dominate another. This means that no matter the sign of *y_2_* + *y*_3_ + … + *y_s_* = 0, ***A**^T^**y*** will have a negative component.

If *y*_2_ + *y*_3_ + … + *y_s_* = 0, suppose that:

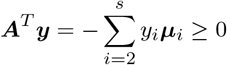

This implies 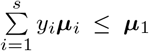 with *y*_1_ = 1, which makes *y*_1_ + *y*_2_ + … + *y_s_* = 1. Here, we cannot have equality since ***μ***_1_,… ***μ**_s_* are linearly independent so we get 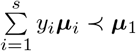. But this contradicts Lemma 1, and we can conclude that non-negative solution ***z*** does exist for the equation ***Az*** = ***b***.

Take the solution ***z*** from above as well as the strictly positive ***y*** invoked from Lemma 3 and define

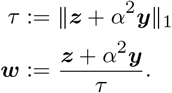

Because of the way that ***z*** and ***y*** were constructed, we have that

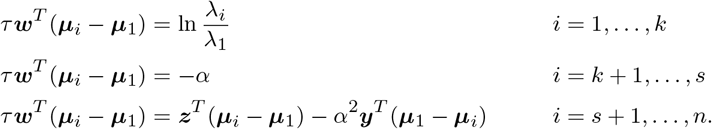

Note that for *i* = *s* + 1,…,*n*,

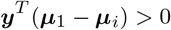

and ***z*** scales at most linearly with *α* so that *α*^2^***y**^T^*(***μ***_1_ − ***μ**_i_*) becomes the dominant term in ***z***^*T*^(***μ***_1_ − ***μ**_i_*) + *#x03B1;*^2^***y**^T^*(***μ***_1_ − ***μ**_i_*) for large enough α. To see this linear scaling, note that ***A*** is of full row rank and our equation is consistent. Using the Moore-Penrose pseudo-inverse ***A***^+^ of the matrix *A*, all solutions to our equation can be expressed as:

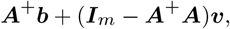

where ***υ*** is an arbitrary vector in 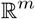. Observe that ***A***^+^***b*** scales linearly with αand that, since ***I***_*m*_ − ***A***^+^***A*** is fixed, we can choose ***υ*** to scale linearly with *α* such that ***A***^+^***b*** + (***I**_m_* − ***A***^+^***A***)*υ* is non-negative, non-zero and scales linearly with *α*.

Since *α* > 0 was arbitrary and 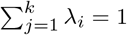, we can let *α* → ∞, resulting in

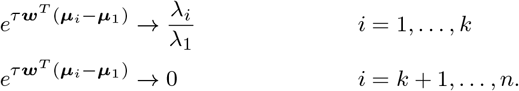

and therefore

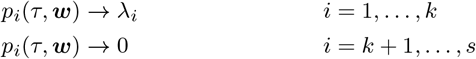

This was the desired result.

*Proof of Proposition 4* Recall Assumption 1 requires that only one vertex of 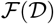 maximizes an individual dimension. Therefore, 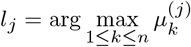 so that 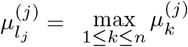, for *j* = 1, 2,…*m*. Notice that as *τ* − ∞:

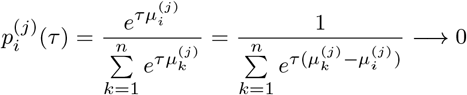

unless *i* = *l_j_*, in which case we obviously have 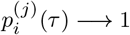. This means that:

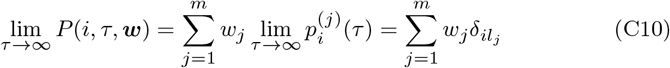

Consequently:

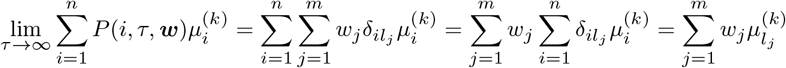

which finally implies:

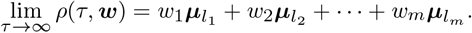

What we have concluded here is that in the limit *τ* → ∞, *ρ*(*τ*, ***w***) is a convex combination of the points ***μ***_*l*__*j*_ for *j* = 1, 2,…*m*. But if we recall the fact that ***μ***_*l*__*j*_. is exactly the point that maximizes objective *j*, then it follows that the image of *ρ*(*τ*, ***w***) on the boundary *τ* → ∞ is precisely the hyperplane Δ.

Furthermore, the selective attention decision-maker with parameters *τ*, ***w*** takes a particularly nice form:

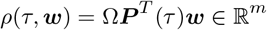

where

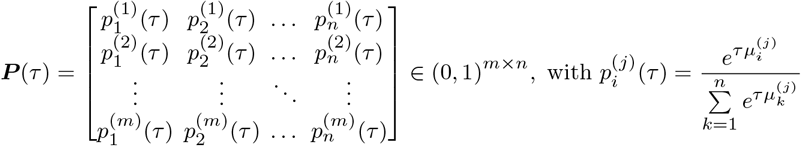

This means that for a given *τ*, *ρ*(*τ*, ***w***) is a linear transformation on 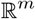 and therefore takes the boundary with *w_j_* = 0 for some *j* ∈ {1, 2,…,*m*} to a boundary point in the image.

## Appendix D Choosing a scale for outcomes

When we analyze AAC tasks, we must assign a numerical value to each outcome. How to do this is not always obvious, especially for harmful outcomes. For example in the spider gambling task (Pittig et al., 2014), one draw of deck *C* might give $25 as a reward and a picture of spider as harm. The natural question is then what numerical value we should assign to the picture of spider?

It turns out that this choice of numerical value does not matter for some of the families of decision-makers. Specifically, we say a family is *scale-invariant* if we were to re-scale any outcome dimension so that the reward structure Ω becomes *A*Ω for some diagonal matrix *A* with positive diagonal entries, then the family’s new feasible region would simply be the original feasible region transformed by A. Since the feasible region for the set 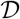 of all decision-makers is just the convex hull of the ***μ**_i_*, then the set of all decision-makers 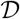 is clearly scale-invariant.

Both the family of softmax decision-makers and the family of epsilon-greedy decision-makers are also scale-invariant. To see this for the former family, suppose that *α_i_* is the ith diagonal entry of *A*. Then when moving from Ω to *A*Ω, we can adjust the (*τ*, ***w***) parameters by introducing 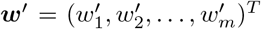 and

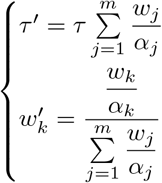

It can be observed that:

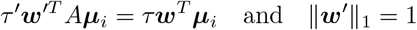

which ensures the probability of choosing action *i* stays the same after scaling. This shows that for every scalarized softmax decision-maker *D* under Ω whose long-term average expected rewards are *ρ*(*D*), there exists a scalarized softmax decision-maker *D*′ under *A*Ω whose long-term average expected rewards are *ρ*(*D*′) = *Aρ*(*D*). Since *A* is invertible, then we can conclude that the family of scalarized soft-max decision-makers are scale-invariant. Furthermore, the same re-scaling of ***w*** to ***w*** also shows that the epsilon-greedy decision-maker is scale-invariant:

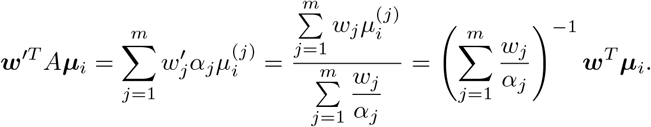

This means that the actions that maximizes the left-hand side also maximizes the right-hand side and vice versa, so the probability distribution of the epsilon-greedy decision-maker remains unaffected.

Interestingly, it is not clear if the family of selective attention decision-makers is scale-invariant. Thus, the choice in scale for outcome dimensions may affect the geometry of the feasible region, beyond a linear transformation, for the family of selective attention decisions.

